# Motif Disruption Domains Lead To Cancer Gene Expression Rewiring

**DOI:** 10.1101/126359

**Authors:** Fabien C. Lamaze, Aurelien Chateigner, Hilary A. Edgington, Marie-Julie Fave, Armande Ang Houle, PCAWG3, Philip Awadalla

**Author notes:** Correspondence to: Fabien C. Lamaze & Philip Awadalla.

## Abstract

Somatic mutations accumulate in non-coding regions of the genome during tumorigenesis, but their functional characterization presents a challenge. Somatic non-coding mutations rarely overlap among patients, which necessitates large sample sizes to detect associations. We analysed somatic mutations called from whole-genome sequencing (WGS) and RNA sequencing (RNAseq) from 3000 tumors across the Pan-Cancer Analysis of Whole Genomes to identify and functionally characterize mutation accumulation and its impact on gene dysregulation in cancer. We identified 1.5 million motif disruption domains (MDDs) across 40 cancer types, which we characterized as pan-cancer targets for recurrent mutation accumulation. These MDDs deregulate gene expression in cancer-specific and pan-cancer patterns by disrupting transcription factor binding sites in regulatory and insulator elements. Disruption is most recurrent across individuals at MDDs in conserved open chromatin, revealing potential drivers. This accumulation of somatic variants targeting regulatory and structural elements in MDDs generates gene expression dysregulation during tumorigenesis.

## Introduction

The discovery of many disease-associated loci in non-coding regions of the genome^1–3^ suggests that these regions are home to a variety of important functions. Recently, evidence has accumulated suggesting a role for non-coding variants in disrupting regulatory elements in cancer, leading to aberrant gene expression and tumorigenesis. Silencing of enhancers and insulators can lead to the dysregulation of epigenetic programs^4^,^5^, notably in colorectal^6^ and prostate cancers^7^. Mutations arising in promoters through the accumulation of somatic single nucleotide variants (SNVs) can affect downstream gene expression^8^, and may even generate driver mutations when arising in certain key promoters such as TERT^8,9^. In addition, SNVs arising in structural elements like insulators can affect gene regulation through the modification of 3D chromatin structure^5,10,11^. However, our understanding of the consequences of SNV accumulation and their clustering in specific regulatory and structural elements of the genome across cancers is incomplete, despite their well documented role in altering gene expression in cancer.

Somatic non-coding variants tend not to be recurrent among patients, which makes the characterization of their functional impact particularly challenging. As a result, large whole-genome sequencing (WGS) and gene expression datasets from paired normal and tumor data are required to capture rare SNVs in order to provide sufficient power to detect their effects on gene regulation. Recent studies have addressed this problem by focusing on regions of SNV accumulation rather than individual SNVs, which greatly increases the power of detection of associations. Two main approaches include the use of functional annotations to identify regions of interest (mainly known regulatory regions), and identifying regions based on clustering of SNVs in the genome across samples. Limiting the scope of a study to known functional regions^2,3,13–15^, allows for greater power, but relies on our limited understanding of the functional role of the non-coding genome^16^. Allowing the data to inform the identification of regions using clustering algorithms^3,17^ prevents the limitations of functional annotation from impacting resulting regions. Here we provide a more comprehensive examination of the impact of SNVs on the gene regulatory program in cancer in several ways. First, whereas previous studies have limited their analyses to either specific mutational targets (e.g. those in previously annotated regulatory elements) or specific targets of dysregulation (e.g. TERT), we used a genome-wide clustering algorithm to identify regions of mutational accumulation, and included all genes as potential targets of dysregulation. In addition to analyzing both WGS and RNA-sequencing data, we incorporate chromatin accessibility data into our analyses in order to identify regions with a higher probability of impacting gene activity. We also take a multi-faceted approach to gene expression regulation by focusing on both regulatory and structural elements as potential targets of mutation.

We performed a comprehensive analysis of SNVs in WGS from paired germline and tumor biopsies of 2835 donors collected from the Pan-Cancer Analysis of Whole Genomes project (PCAWG), as well as RNAseq data from 1188 donors including 121 paired tumor and normal tissue biopsies from the same individuals. Here we show that while the vast majority of non-coding SNVs are observed at low frequency across individuals, the majority cluster in recurrent regions of high mutation density across cancers, which we call motif disruption domains (MDDs). Analyzing MDDs enabled us to uncover novel pan-cancer associations between motif disruption and gene dysregulation, which the analysis of non-recurrent individual SNVs would have made undetectable. Additionally, we discovered extremely recurrent disruptions at MDDs we classify as potential novel driver domains, some of which colocalize with known cancer regulators such as MYC and HMGIY.

## Results

### Motif disruption domains colocalize with regulatory and structural elements

The vast majority of SNVs in the PCAWG data are non-recurrent (96.8% singletons), as previously observed^16,17^. In order to detect clustering of SNVs across the genome, we concatenated continuous genomic areas encompassing at least two SNVs no more than 100bp apart across all tumor samples. 100bp was found to be the distance at which we no longer saw a substantial change in the number of MDDs identified when we tested a range of distances from 25bp to 1600bp. This approach allowed us to identify 1,507,105 pan-cancer densely mutated genomic domains of median size 1.2kb, which we will refer to as motif disruption domains (MDDs). Among these MDDs were previously reported highly mutated non-coding regions, such as the TERT promoter^9^ (MDD at chr5:1295032-1295670). Despite the non-recurrence of individual SNVs, 53.7% of all SNVs fall into an MDD, which tend to accumulate outside of genes, primarily in distal intergenic regions (Fig. 1a), at a median distance of 21,395bp to a TSS. MDDs fall primarily in heterochromatin (19-32% within each cancer type) and quiescent DNA (53-72%), concordant with the accumulation of SNVs in DNAse free regions^18^. However, 10% to 22% of MDDs overlap with active transcription start sites (TSS) or enhancers (Fig. 1b). These results show that while individual SNVs may not be recurrent among individual tumours, they fall into regions of high SNV density that may have functional importance during tumorigenesis, and using MDDs as the unit of analysis increases our power of detecting those functional associations.

**Figure 1.**
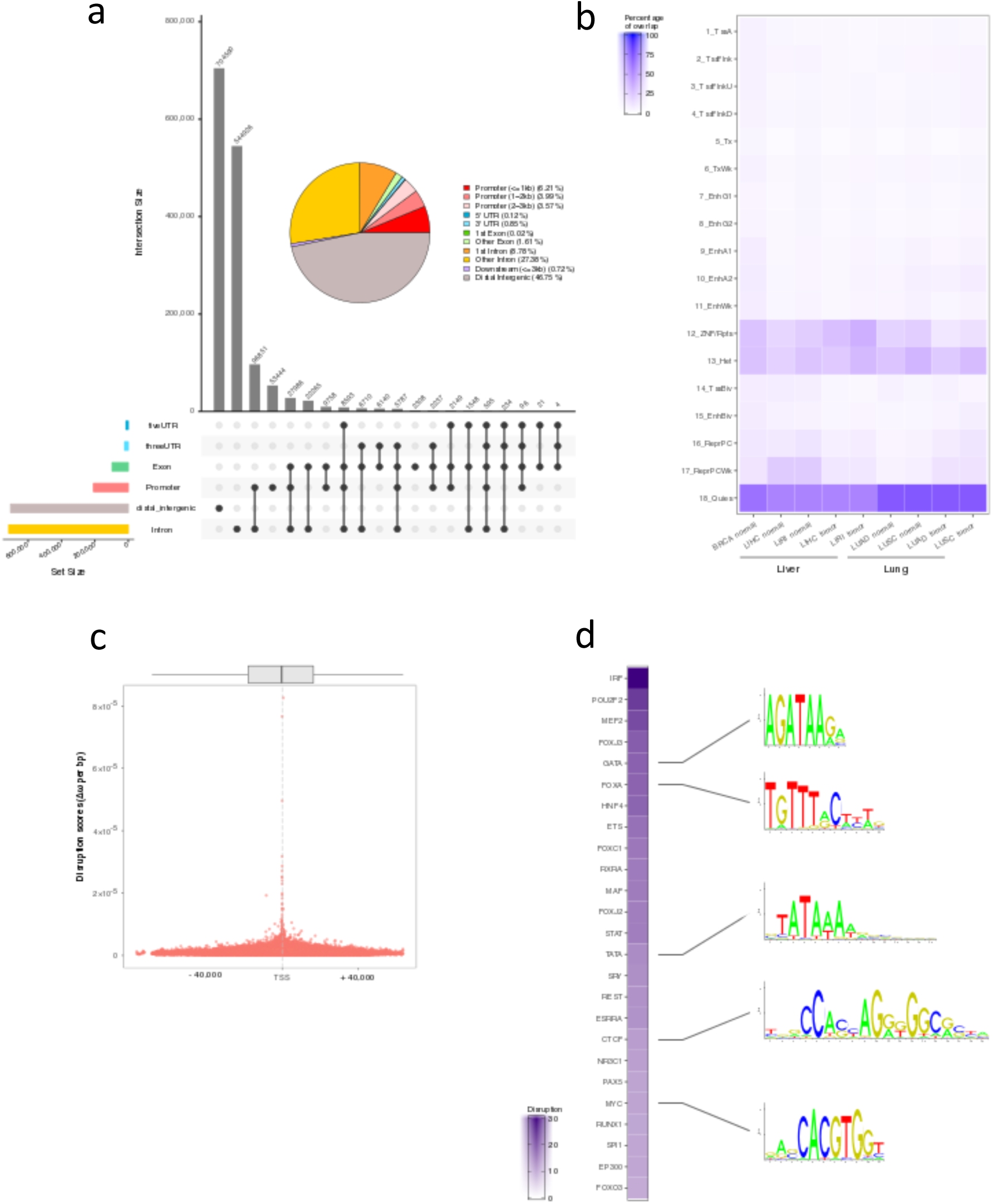
MDD genomic landscape colocalize with regulatory and structural elements. **(a)** Genomic annotation of MDD using GENCODE v19. (**b**) Heatmap showing the percentage of overlap between MDDs containing disruption scores > 0 and the functional genome. The ChromHMM 18-state model in normal and tumor cells (see Methods) was used to characterize the functional genome. (**c**) MDD disruption scores compared to distance from a TSS. (**d**) Top 25 most disrupted motifs in the 1% most disrupted MDDs

The non-coding genome is the target of almost 1400 transcription factors (TFs)^19^ that contribute to the regulation of transcription of nearby genes by binding to specific motifs. To investigate the TF binding motifs in MDDs that have been targeted by SNVs, we computed a motif disruption score for each TF motif. The scores for all motifs within each MDD were summarized to produce a “MDD disruption score”. MDD disruption scores were negatively correlated with distance to a TSS (*p* < 2 × 10^−16^, □ = -0.10; Fig. 1c), supporting their potential role in the regulation of gene expression. Interestingly, MDDs within the top 1% of motif disruption scores capture binding sites of TFs and cofactors known to bind to promoter or enhancer regions (Fig. 1d; Supplementary Table 1). Those associated with promoters include binding sites of general TFs, core elements of the proximal promoter (TATA box), and known proto-oncogenes such as MYC, STAT, SPI1 from the E26 transformation-specific family, and those associated with enhancers include binding sites of pioneer TFs (FOXA and GATA) and chromatin remodeler binding sites (EP300). We also found that CTCF (CCCTC-binding factor) binding sites were included in the top 25 most drastically disrupted motifs across cancer (Fig. 1d). SNV accumulation at binding sites of CTCF ^5,10,11^, a boundary element that recruits cohesin to stabilize the chromatin loop^12^, confirms the frequent disruption through mutation of CTCF in various cancers^5,11^. The relationship between disruption at an MDD and colocalization with regulatory elements critical to proper gene regulation points toward a disruptive role of MDDs in the gene expression program though structural and/or regulatory element disruption, likely to affect long-range enhancer-promoter interactions.

MDDs may alter transcription through two main processes in cancer: MDDs overlapping with regulatory regions disrupt TF binding sites, resulting in the alteration of the normal gene expression program of the cell, and MDDs at chromosome structural elements impact the integrity of larger cohesin-mediated loops called insulated neighborhoods and topologically associated domains (TAD). We focus specifically on the structural element CTCF, as it has been previously shown to be the target of mutation accumulation in cancer^5,10,11,15^. Moreover, somatic mutations impact the binding affinity of CTCF^20^, and deletions overlapping CTCF binding sites in T-cell acute lymphoblastic leukemia was associated with aberrant expression of TAL1 and LMO2 genes^5,10,11^. However, a relationship between mutation accumulation at CTCF and aberrant gene activity in the surrounding neighborhood has not previously been shown. By examining associations between TF motif disruption at MDDs and the gene activity of neighbor genes in the context of overlap with regulatory elements we can identify MDDs that are important to the gene regulatory program during tumorigenesis.

### MDDs disrupt the cell gene expression program

To test whether MDDs impact TF binding sites or chromosomal structural elements within and among cancer types, we first investigated the MDDs located in accessible regions of the genome, as assessed by their enrichment for DNase1 in 7 cancer types for which we have paired normal and tumor gene expression data. We looked for associations between motif disruption scores and changes in gene expression between tumour and normal samples using Coinertia Analyses (COIA)^21^. 33,485 MDDs contained regulatory elements overlapping accessible DNA. The pattern of DNase1 enrichment at MDDs is generally tissue specific, with a maximum of 64.8% overlap at an MDD among all tumour and normal tissues (Fig. 2c, Figure S1). COIA of each cancer type revealed a number of associations between differential gene expression between normal and tumor biopsies and the disruption scores of MDDs overlapping DNase1 (Fig. 2c), suggesting that MDDs influence the deregulation of gene expression between normal and tumor tissue in accessible chromatin regions. Second, we investigated MDDs that overlap with neighborhood domains identified using chromatin interaction analysis by paired-end tag sequencing (ChIA-PET) for SMC1, a subunit of cohesin^22^ in the 7 cancer types for which we have paired normal and tumor gene expression data. Similarly, we observed significant associations between CTCF disruptions in the 746 MDDs overlapping these elements and gene dysregulation within the topologically associated domain of that neighborhood (Fig. 2d). In both cases we observe a cancer-specific pattern of association between MDD disruption and gene expression alteration (Fig. 2c, 2d), suggesting that tissue-specific gene regulation in the cell of origin is a predictor of the dysregulation in gene expression during tumorigenesis. While the interaction network between MDDs and gene expression differs among tumors, a recurrent analysis of correlation coefficients from gene-MDD associations reveals a commonality of dysregulation associated with specific MDDs among cancers (Fig. 2c, 2d).

**Figure 2.**
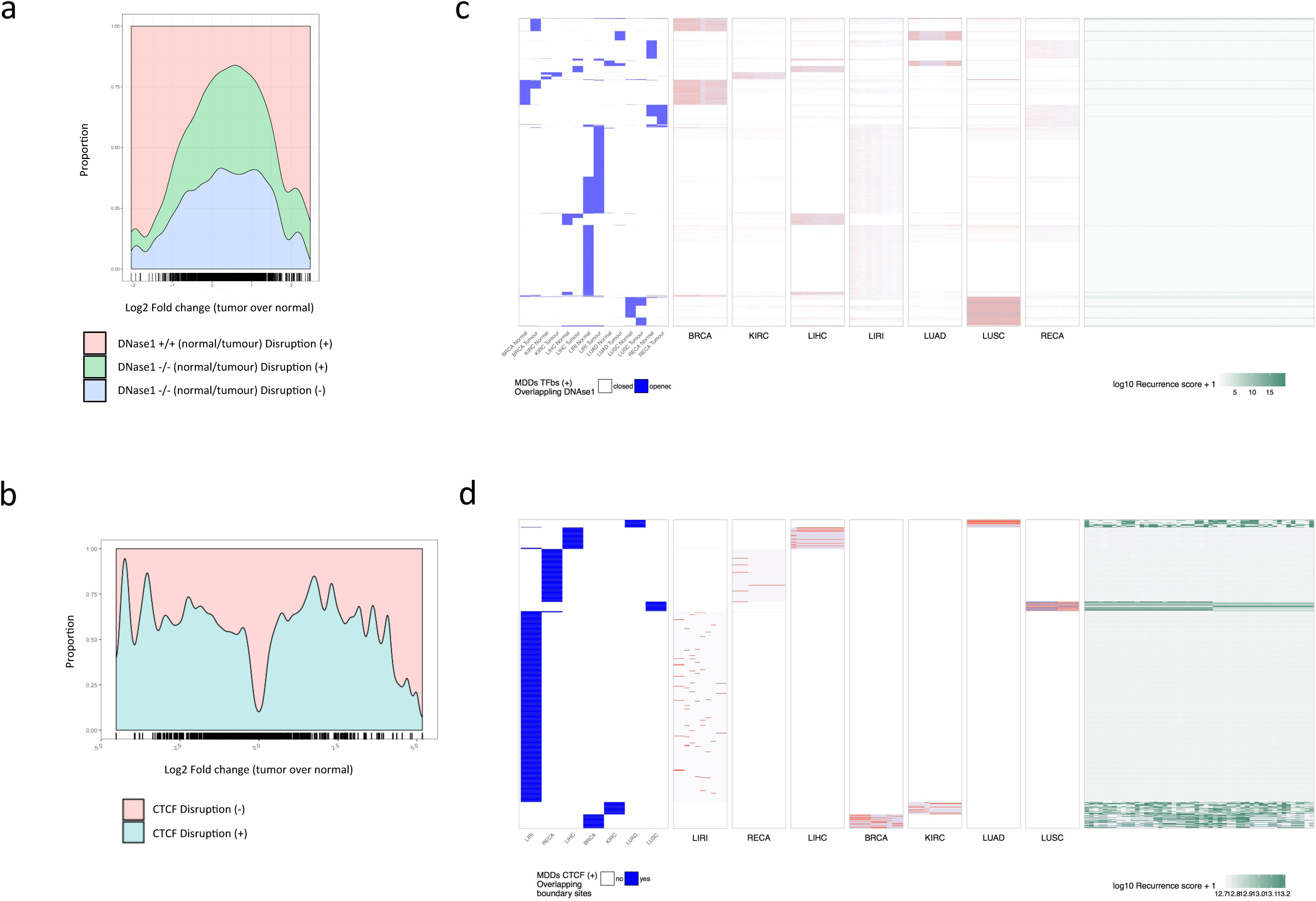
MDDs disrupt regulatory and insulator elements leading to gene dysregulation. (**a**) Coding genes having their promoter in open chromatin in normal and tumor tissue and overlapping an MDD with disrupted TFBSs are over represented in extremely dysregulated genes. (**b**) Genes having the boundaries of their TAD disrupted are over represented in the extreme fold change values (**c**) The correlation between MDD disruption and gene expression alteration in regions overlapping DNase1. The left panel represents the overlap between the 33,485 MDDs (in rows) and the DNase1 regions (blue = overlap, white = no overlap), specifically computed for the tumor and the associated normal tissue of each cancer (in column). Euclidean distance was used to cluster the columns and rows. Row order is consistent among all panels. The 7 middle panels represent the correlation between the disruption scores for the same MDDs (in rows) and fold-change in gene expression for the 11,243 genes with differential expression between tumour and normal pairs. Only the 10 genes with the highest correlations are shown for each cancer. All panels are on the same scale, from -1 to 1. The top 100 correlations for each cancer are shown in table S4. Finally, the right panel represents the logarithm of the recurrence score (+1) of each correlation between a MDD and a difference in gene expression across the 7 cancers. A higher log correlation score means that there is recurrent disruption-dysregulation correlation (negative or positive) among the 7 cancers. (**d**) Correlation between disruption at MDDs overlapping CTCF and TAD boundary elements and neighborhood gene dysregulation. The left panel represents the overlap between the 746 MDDs (in rows) and the CTCF and TAD regions (blue = overlap, white = no overlap), specifically computed for the tumor and the associated normal tissue of each cancer (in columns). As in c, the columns and rows are clustered by euclidean distance and the row clustering is consistent across the other panels. The 7 middle panels again represent correlations between the MDDs (in rows) and fold-change in gene expression for the 20,801 genes found in insulated neighborhoods. Again, only the 10 genes with the highest correlations are shown in each panel and all are on the same scale (−1 to +1). The right panel, as in c, represents the recurrence of a correlation between disruption and gene expression alteration at an MDD across the 7 cancers.

### MDDs impact important biological functions in cancer

An interesting result from the COIAs is the frequent number of associations we observed between disruption in MDDs and dysregulation of non-coding RNAs. For example, we found that disruption led to dysregulation in the microRNA miR-187, which has previously been associated with tumorigenesis in several cancers (e.g. prostate^24^, nasopharyngeal^25^, renal^26^, pancreatic^27^, thyroid^28^, breast^29^, and oesophageal^30^) and is known to increase tumor invasiveness^29^, as well as miR-8, which also was shown to promote tumorigenesis^31^,^32^. We also observed associations between MDD disruption and dysregulation of small nucleolar RNAs (snoRNAs), which perform housekeeping functions by regulating translation of other RNAs and can also represent cancer diagnostic/prognostic markers (e.g. SNORD89). snoRNAs are regulated by the MYC transcription factor, which are found to be highly disrupted and associated with dysregulation of gene expression.

We characterized the 300 genes with the strongest correlations with MDD disruptions from each COIA using molecular interaction networks to visualize interactions between the genes and their regulators (Supplementary Tables 2a and 2b). Although the COIAs included both protein-forming as well as non-translated genes, we only included protein-forming genes in the networks. In addition to known associations with cancer among the genes impacted by disruption at an MDD enriched with DNase1, including TRIM23, SAA1, LTBR, CHEK2, and SMAD3, we observed a number of interactions between our query genes and cancer-associated genes or TFs (e.g. MYC, CDK4, EP300, FOXO1, and RHOA) (Fig. 3a). Though we found fewer correlations between disruptions at MDDs overlapping with CTCF binding sites and gene expression changes than MDDs enriched for DNase1, they also included genes with known associations with cancer (e.g. TSPAN1, DES, and SFTPA) as well as interactions with known cancer regulators (e.g. SOX2, EP300, CD14) (Fig. 3b.). Interestingly, there was substantial overlap between connecting nodes in the network and transcription factors we found to be highly disrupted (Fig. 1d.), including JUN, EP300, STAT3, NGF4A, and MYC in the DNase1 network (Fig. 3a.), and EP300, SOX2, and SP1 in the CTCF network (Fig. 3b.). This suggests that these factors are not only highly disrupted among individuals, but that they are integral to the dysregulation of gene expression during tumorigenesis.

**Figure 3.**
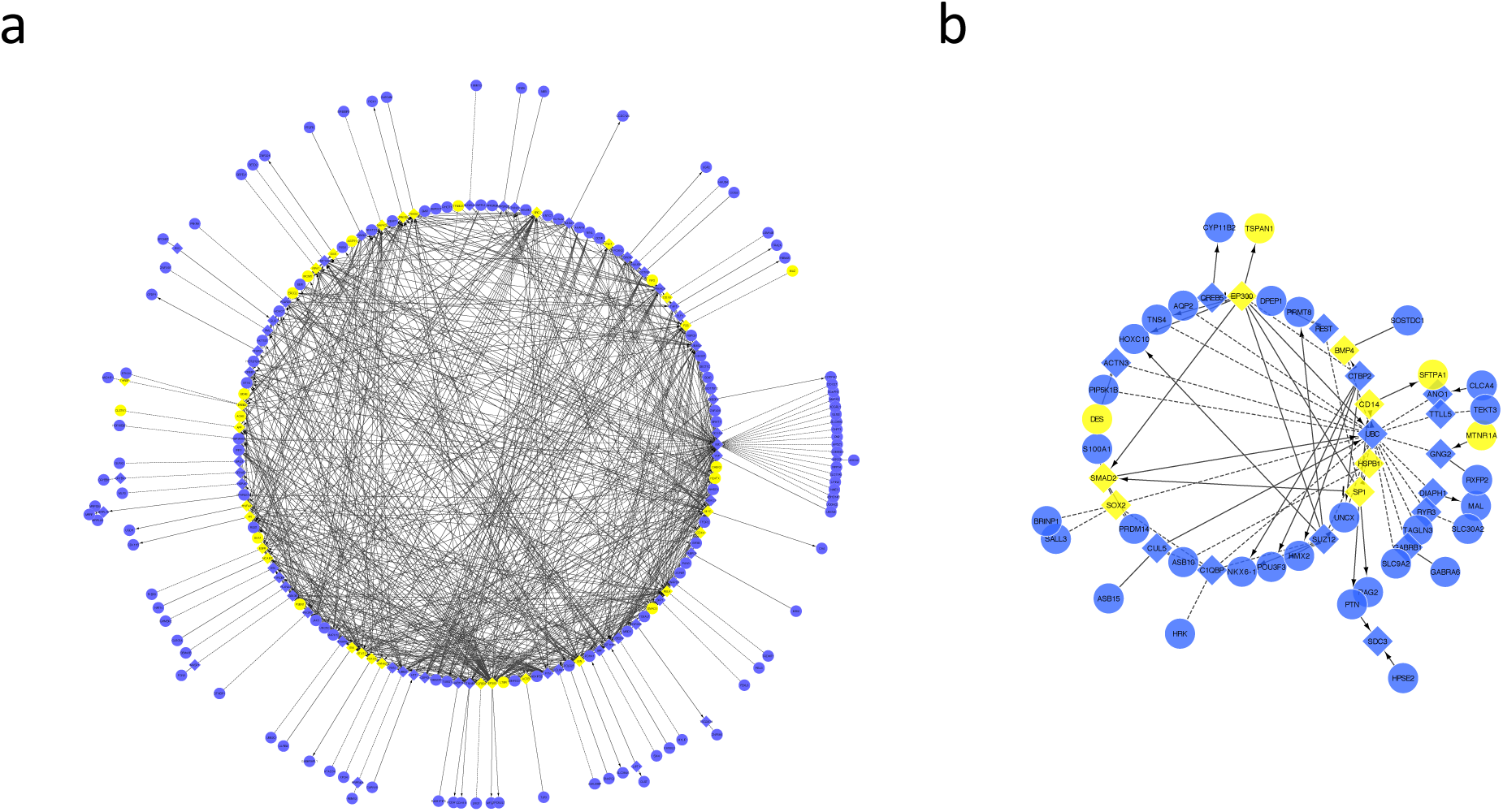
Disrupted transcription factors within MDDs interactant with dysregulated genes. Recurrently dysregulated coding genes associated with MDD disruption (see Fig. 2c and 2d) are represented by circles alongside with their regulators (diamonds) in a network constructed using Reactome data, including the 300 strongest correlations resulting from coinertia analyses. Two networks were constructed, one using genes associated with disruptions in MDDs enriched for DNase1 (**a**), and one with genes associated with disruptions in MDDs overlapping CTCF and TAD (**b**). Genes and regulators involved in tumorigenesis are highlighted in yellow.

### MDDs enable the identification of potential non-coding cancer-specific and pan-cancer drivers

For an MDD to contribute to the regulation of cancer progression, the chromatin (i) should be in an open state in the normal cell of origin and (ii) should not become inactivated during clonal range expansion. To identify potential regulatory driver MDDs, we studied motif disruption frequency in MDDs grouped by chromatin accessibility, as we expect disruptions in regions that are important to cancer development to be more recurrent. MDDs were grouped in four categories based on DNAse enrichment in different tissue types: enrichment in normal and tumor tissues, in normal tissue only, in tumor tissue only, and in neither normal nor tumor. We first tested whether disruption recurrence differs among DNase1 categories across all MDDs, and found that while the vast majority of MDDs were disrupted in very few individuals, the recurrence of disruptions across individuals within cancers was significantly different among DNase1 categories and cancers, with a significant interaction effect (p<0.0001 for all comparisons; for full model results see Supplementary Table S3a). The increased recurrence among MDDs in open chromatin in both tumor and normal tissue suggests that we have captured potential drivers in this group.

To examine potential driver MDDs more specifically, we focused on the outliers of the distribution of proportion of individuals with a disruption within each cancer and DNase1 category, i.e. the MDDs that are ‘extremely recurrent’. We observed a pattern whereby, as with the full dataset, outlier MDDs are more frequently found across many samples when found in regions accessible in both tumour and normal tissues (Fig. 4a), with the highest recurrence (100%) being observed for an MDD among cutaneous melanoma patients (Fig. 4b). This pattern is observed across cancer types, and recurrence is significant among DNase1 categories, cancers, and the interaction between them (p<0.0001 for all comparisons; for full model results see Supplementary Table S3b). We consider ‘extremely recurrent’ MDD disruptions in regions of open chromatin in both tumor and normal tissue to represent potential non-coding cancer-specific drivers (Fig. 4a; Supplementary Table S4). We identified gene ontology groups, biological pathways, regulatory motifs, and proteins statistically enriched in the outlier MDDs for each cancer (for full enrichment results, see Table S5a). For example, outlier MDDs in breast cancer were enriched for ETF, a transcription factor associated with overexpression of the oncogene EGFR^33^, and E2F1, which has a crucial role in the control of the cell cycle and regulation of tumor suppressors, as well as being a target for DNA tumor viruses^34^. Furthermore, colorectal adenocarcinoma had a large number of significantly enriched transcription factors, including HOXB5, a known contributor to several cancers^35,36^, NFATC2, which is involved in cancer immune dysregulation^37^, Egr1, a regulator of multiple tumor suppressors^38^, and HMGIY, an architectural transcription factor associated with neoplastic transformation and metastatic progression^39^. In most cancers we found enrichments of several homeobox binding sites (e.g. DLX1, GSX2, CRX, LMX1A, Ipf1, Pitx3, LMX1B, and NKX6-1) as well as other factors of importance such as TBP and OC-2, and GATA, and enrichments of gene ontology groups involved in important cell processes such as cell-cell adhesion, localization, and communication.

**Figure 4.**
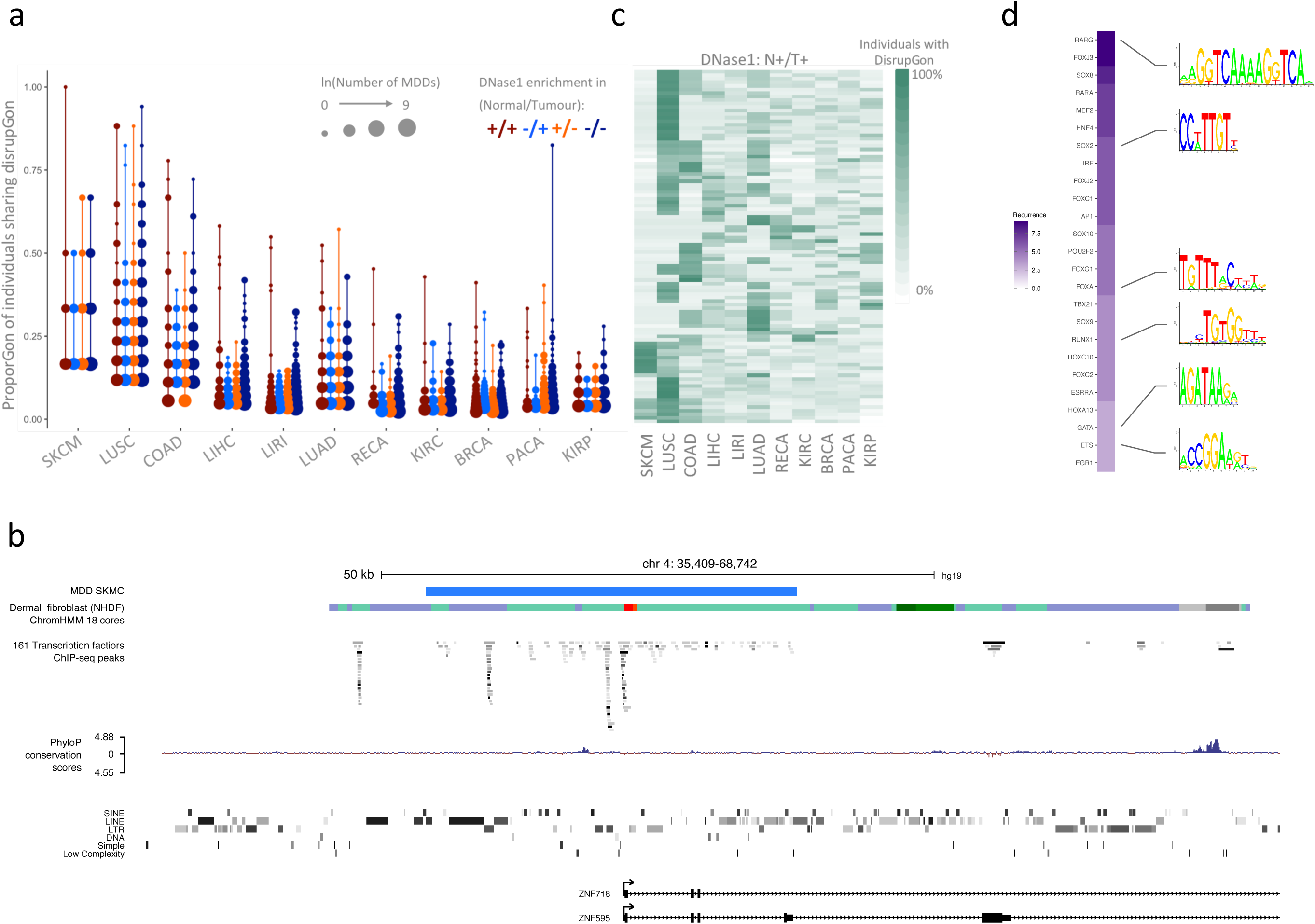
Extremely recurrent MDDs are potential cancer divers. The proportion of individuals sharing a disruption at a particular MDD in each of four DNase1 enrichment categories were compared for each of eleven cancers. (**a**) While the vast majority of MDDs have a low proportion of disruptions shared among individuals, visualization of the outliers of the distribution of proportions reveals that for the majority of cancers, MDDs in open regions for both tumour and normal samples contain disruptions in a higher proportion of individuals than any other DNase1 category. (**b**) A profile of the genomic, chromatin, and ChiP-seq binding landscape underlying the top outlier in skin cutaneous melanoma (SKCM) from Fig. 4a. (**c**) Commonality of disruptions in these accessible domains suggests that these may include drivers that are critical to cancer genesis and maintenance. The proportion of individuals with disruptions in each tumour-open and normal-open MDD was compared across cancers with a recurrence analysis in order to identify commonly disrupted pan-cancer MDDs. (**d**) While disruption sharing across cancers was minimal, there was considerable overlap among the specific TF motifs where the most disruptions occurred among the cancers. Here we show the most frequently disrupted motifs across cancers (for individual cancers see Supplementary Fig. S2).

We used a recurrence analysis to identify MDDs with high frequencies of disruptions across all cancers, potential pan-cancer drivers, by ranking MDDs by the proportion of individuals with a disruption for each cancer, then taking the median rank for all cancers for each MDD (Fig. 4c). We found that regions associated with meiosis and meiotic regulation, recombination processes, and the regulation of RNA Polymerase I, as well as for known cancer regulators such as POU4F1^40^, ETF^33^, and Oct-4^41^ were among the processes and factors significantly enriched in the top 500 pan-cancer recurrently disrupted MDDs (for full enrichment results, see Table S5b).

To further characterize potential driver MDDs, we determined which transcription factor motifs were recurrently disrupted among them. We identified all TF motifs in outlier MDDs in regions of conserved open chromatin, and extracted the most frequently disrupted motifs within each cancer (Supplementary Fig. S2). The 25 most recurrently disrupted TF motifs across cancers were identified based on the number of times a TF motif was observed among the most disrupted TF motifs within each cancer (Fig. 4d). TFs in these most recurrently disrupted motifs overlapped with those with the highest disruption scores (Fig. 1d), as well as those associated with gene dysregulation in the COIAs (e.g. JUN, FOXO1, MYK, SOX2, and SP1). Among the most recurrently disrupted TF motifs are other factors of note such as ETS^42^, HNF4^43^, and RUNX1^44^. We conclude that these MDDs may be classified as potential drivers, as they represent SNV targets of considerable importance to gene expression alterations involved in cancer.

Next, to document further the roles of MDDs in tumorigenesis, we tested whether the strength of motif disruption or the motif disruption recurrence is a stronger predictor of alterations in gene expression. We used a linear model to test the relationship between log-fold changes in gene expression and (1) mean disruption score at the associated MDD, (2) the proportion of individuals exhibiting disruption at the MDD, and (3) the interaction between the two. We found that the model was significant, but explained very little of the variance (p = 0.0268, R2 = 1.044×10^−5^). Surprisingly, while there was a slight but significant relationship between change in expression and the proportion of individuals with a disruption (p = 0.027), there was no significant relationship between the change in gene expression and average disruption score (p = 0.348; for full model results see Supplementary Table 6). Collectively, these results suggest that the disruption of regulatory elements may have a threshold rather than an additive effect, where after a disruption occurs the alteration in gene expression is not further exacerbated by additional disruptions.

## Discussion

In this work we present a novel method of characterizing the functional impact of noncoding somatic mutations. By integrating whole-genome sequencing, RNA sequencing, and chromatin accessibility data, we have developed a new classifier of non-coding mutation accumulations that disrupt regulatory and insulator elements, and present a comprehensive view of how mutation accumulation impacts gene expression in cancer. By identifying MDDs that are recurrent targets of mutation across all cancer types, and using these MDDs as units of analysis, we had sufficient power to detect associations between mutation accumulation and genome-wide gene expression dysregulation despite the non-recurrence of SNVs. We found that the determined regions of recurrence are targets of mutation accumulation that have direct impact on the regulation of gene expression both through regulatory and structural elements. The targeted transcription factor binding motifs at these MDDs include a wide array of well documented gene regulators such as TBP and structural elements like the insulator CTCF, the latter known to be a target of noncoding SNVs in colorectal and other cancers^5,11^. The deletion of the entire CTCF binding site is known to lead to aberrant gene activity^5,11^, and here we show that the accumulation of point mutations leads to similar dysregulation. This suggests a mechanism to explain the known contribution of dysfunction in topologically associated domains to cancer initiation and progression, and to other diseases^45^.

Our analysis of MDDs revealed potential cancer-specific and pan-cancer drivers through the examination of recurrently disrupted MDDs in regions of open chromatin. We found enrichments in these putative driver MDDs of both known cancer regulators such as HOXB5 and Egr-1 as well as biological processes and transcription factors without known associations with cancer. Our approach highlights the large benefit of considering recurrently mutated genomic regions, overpowering that of relying on individuals SNVs, to expose the role of non-coding somatic mutations in gene dysregulation in cancer. Interestingly, our comparison of the strength of disruption at an MDD, the recurrence of those disruptions, and their effect on gene dysregulation suggests that there is a threshold effect to gene dysregulation, where increasingly strong disruptions at an MDD do not contribute additively to the alteration in gene expression. Our results suggest that one or a small number of non-coding somatic mutations can cause aberrant gene activity leading to tumorigenesis, and that some of these regions of mutation accumulation are also recurrent targets across cancers, potentially having a role in driving the cancer phenotype. Altogether, our identification of motif disruption domains revealed targets of somatic non-coding mutation accumulation that play a key role in the alteration of gene expression during cancer development, and which include potential novel drivers of tumorigenesis.

## Methods

### Whole genome sequencing and somatic point mutation calling

The PCAWG project sequenced the genomes of tumor biopsies and germline normal tissue (whole blood) with high quality standards for 2588 donors (freeze v1.4; syn5871617). Variants specific to the tumor were called excluding tissue-specific and individual-specific germline variants from all tumor variants by PCAWG working group 1. From the 2588 white-listed tumor biopsies initially available, 959 had whole genome and RNA sequencing available for the same tumor, and 300 of these had RNAseq data available for the adjacent normal tissue.

### Mutational domains

To determine the boundaries of the mutational domains, we used the bedtools merge function from bedtools^49^ (version 2.24.0). All single nucleotide variants (SNVs) from the 2588 whitelisted tumor biopsies available, falling within 100 bp of each other were clustered together in mutational domains. The 100bp distance between clustering SNVs was chosen by systematically identifying domains using different distances ranging from 25bp to 800bp, then choosing the distance where we observed a diminishing rate of change on the asymptotic curve of the number of domains identified. As some recurrent SNVs are neighbour-less, and would comprise 100% of 1bp regions, giving undue weight to individual SNVs, all regions were extended by flanking 50bp, to be at least 101bp in length. These domains were discarded from subsequent analyses if unique to a single donor.

### Motif search and motif disruption scores

A transcription factor binding site (TFBS) search was done using the R^50^ package motifbreakR^51^ (version 1.2.2) for each individual and each single nucleotide alteration, we searched for TFBS affected by the variant, and computed disruption scores for each TFBS. The score of each motif was computed by comparison with the position probability matrices from the reference genome BSgenome.Hsapiens.UCSC.hg19 (version 1.40.1). The germline score was calculated using the SNVs called from the germline WGS sequences. Finally, the somatic score was calculated based on the somatic SNVs called from the tumor WGS sequences. The motif disruption or relative entropy □ was computed as per Equation 1, where i = 1,…n, M is a frequency matrix of width n (length of the motif sequence), and b_i_ (set to 0.25, an approximation of the human background frequency ^52^) is the background frequency of the letter i:

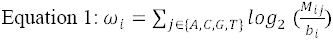

Total motif disruption scores for each donor were computed using distance matrices that summarise the effects of somatic SNVs on all motifs when compared with the reference motifs from the germline sample of the same donor. Owing to the variation among conservation scores within TFBS motifs, we computed the percentage of disruption scores for the motif for the reference allele (WGS of the germline) and the somatic SNV, defined as:

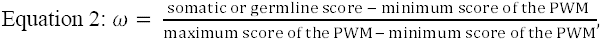
where PWM is the position weight matrix.

We then summed the absolute difference between the germline and somatic percentages of motif disruption (□□) where multiple TF binding motifs overlap at a given single nucleotide alteration locus, and corrected for the size of the region to determine the score of the disruption within a given region for each donor.

### Most targeted motif in motif disruption domains (MDDs)

MDDs in the top percentile of overall motif disruption scores were used to select the most targeted TFBS. We merged motifs associated with the same TF to avoid overestimation of redundant TFBS disruption scores at a same loci by taking the lowest disruption score for this motif. The scores of different TFBSpresent at the same loci were summed. Finally, only the scores of strongly disrupted motifs (as defined by motifbreakR ^51^) were selected in these MDDs.

### Annotation of the motif disruption domains and ChIPseq processing

Gencode v19 was used as the reference for the gene and promoter annotation (Fig. 1a). Cancer-specific chromatin state data were retrieved from the Epigenomic Roadmap consortium, and TF binding site locations in A549 (lung) and HepG2 (liver) cancer cell lines were extracted from ENCODE narrow peaks files. Chromatin state within each MDD was quantified by computing the ratio of nucleotides exhibiting each chromatin state. To functionally annotate the MDDs we used DNase1 peaks from the ENCODE and Roadmap consortia (see Table 1) as well as SMC1 ChIA-pet neighborhood domains (GSE68977). To assess if MDDs fall within potentially active regulatory regions, we computed the overlap between MDDs with DNase1 regions within tissue and cancer types. We considered MDDs to cause structural chromatin alteration only if they contained a disruption of CTCF binding motif, determined by computing the overlap between MDDs and SMC1 binding domains.

### RNAseq data preprocessing

After excluding RNAseq data flagged for quality issues, our RNAseq dataset included 300 normal and tumor paired biopsies from freeze v1.4 for which we also had WGS data, processed by the PCAWG consortium groups 3 and 14 (syn5871617). Raw sequencing FASTQ files were aligned using STAR and TOPhat aligners, and read counts per gene were calculated for both the STAR and TOPHAT pipeline using htseq-count with the PCAWG reference GTF, a modified version of Gencode v19 (syn3221170). We filtered out mitochondrial genes, and genes for which 90% of the donor had FPKMuq values under 10. We used the SVA package to estimate one surrogate variable while controlling for the cancer type and tumor or normal status for analyses using both WGS and RNAseq data, and only for cancer type for analyses using only WGS data, followed by a GLM with a poisson distribution to regress out the surrogate batch effect. The residuals from the regression analysis were used to compute the fold change in expression between paired tumor and normal tissue biopsies in analyses using WGS and RNAseq data.

### Coinertia analyses

Coinertia analyses were performed using the R packages *ade4^53^* (version 3.3.0). The gene expression and disruption matrices were normalized to one unit variance and centered prior to running a principal component analysis (PCA). PCA was performed (center=TRUE, scale=TRUE, scannf=F, nf=2) on a matrix containing the total individual disruption scores calculated previously for each cancer type, and another was performed on the gene expression fold change matrix. Then, we performed a coinertia analysis between the two PCAs (scannf = FALSE, nf=2). The coinertia analysis resulted in cross tabulated correlation coefficients between the normalized sum of the disruption scores (□□) and the batch corrected fold change of expression of a gene. Then, we computed a Monte-Carlo Test on the sum of eigenvalues of a co-inertia analysis based on RV coefficient with permutation to assess the overall correlation between the gene expression and disruption matrices. We extracted the 300 coding genes with the top correlations with an MDD disruption score for the DNase1 and CTCF COIAs, and visualized their interactions with molecular interaction networks using the Reactome database (2015) in Cytoscape^23^. We included linkers in the visualization to examine regulators of the genes.

### Disruption recurrence analyses

Data for the disruption recurrence analyses was processed in R version 3.3.2. We first used the packages *GenomicFeatures^54^, GenomicRanges^54^, rtracklayer^55^, data.table^56^, reshape2^57^*, and *tidyr^58^*. For each cancer type, we used ChIP-seq data (detailed previously) to determine whether each MDD fell into a region of DNase1 enrichment for both the tumour and normal state. These designations became our chromatin accessibility categories (DNase1 enrichment-Tumour/DNase1 enrichment-Normal; DNase1 enrichment-Tumour/No DNase1 enrichment-Normal; No DNase1 enrichment-Tumour/DNase1 enrichment-Normal; No DNase1 enrichment-Tumour/No DNase1 enrichment-Normal). We then transformed our MDD disruption scores into a binary score (disrupted or not disrupted) for each individual at each MDD, and calculated the proportion of individuals with a disruption at each MDD for each cancer type. We used an Analysis of Variance (ANOVA) to compare the proportions of individuals with a disruption among DNase1 categories, including cancer type in our model to control for inter-cancer differences in disruption recurrence. As we are primarily interested in identifying drivers of tumorigenesis, and therefore in the most shared disruptions across individuals, we used the scores() function from the R package *outliers^59^* to extract for each cancer and accessibility category the observations above the 99th percentile, based on normalized z scores. We tested for differences among the proportion of individuals with disruptions among each accessibility and cancer type category in these outlier MDDs using ANOVA, and visualized the data using *ggplot2^60^*.

We next used the MDDs falling in the DNase1 enrichment-Tumour/DNase1 enrichment-Normal category to identify those with the highest pan-cancer recurrence. We only included MDDs with disruptions seen in at least 50% of cancers. For each cancer, we ranked all MDDs based on the proportion of individuals with a disruption, then took the median ranking of all cancers for each MDD. This median value represents the combination of frequency of disruptions at an MDD both within a cancer and among cancers. We visualized the top one hundred most recurrently disrupted MDDs across cancers using a heatmap of the proportion of individuals with disruptions for each cancer.

To characterize putative driver MDDs identified by the recurrence of disruptions within and among cancers in regions of conserved open chromatin, we used the web-based software program g:Profiler^61^ to search for statistical enrichment of gene ontology groups, biological pathways from KEGG and Reactome, and regulatory motifs from TRANSAC and miRBase. Enrichment analysis was performed for each cancer-specific group of outlier MDDs and among the 500 most dysregulated pan-cancer recurrent MDDs.

In order to characterize the relationship between the recurrence of a disruption among individuals and the strength of that disruption (represented by the MDD’s disruption score), we calculated the average disruption score of an MDD for each cancer, excluding data from individuals with no disruption. After visualizing the data, we tested for a relationship between the log-transformed average disruption scores and the proportion of individuals with a disruption using a linear regression. We then tested for relationships between alterations in gene expression and both the average disruption score and the proportion of individuals with a disruption at an MDD, along with the interaction between the two, using a linear regression.

## Abbreviations

MDD: Motif Disruption Domain
PCAWG: the Pan-Cancer Analysis of Whole Genomes project
SNV: Somatic single Nucleotide Variants
TFBS: Transcription Factor Binding Site

## Author Contributions

Conceived and designed the experiments: FCL and AC. Analyzed the data: FCL, AC, and HAE. Generated the data: PCAWG. Wrote the paper: FCL, AC, HAE, MJF, AAH and PA.

## Supplemental Information

Supplementary Table 1: Ordered list of the MDDs with the top 1% highest disruption scores.

Supplementary Tables 2a and 2b: Genes most recurrently associated to a high MDD disruption overlapping DNase1 or CTCF and TADs across the 7 cancers studied.

Supplementary Tables 3a and 3b: Full models.

Supplementary Table 4: MDDs that are ‘extremely recurrent’.

Supplementary Table 4: Top 100 correlations between MDD disruption and gene expression difference for each cancer.

